# Sex differences in brain correlates of STEM anxiety

**DOI:** 10.1101/528075

**Authors:** Ariel A. Gonzalez, Katherine L. Bottenhorn, Jessica E. Bartley, Timothy Hayes, Michael C. Riedel, Taylor Salo, Elsa I. Bravo, Rosalie Odean, Alina Nazareth, Robert W. Laird, Matthew T. Sutherland, Eric Brewe, Shannon M. Pruden, Angela R. Laird

## Abstract

Anxiety is known to dysregulate the salience, default mode, and central executive networks of the human brain, yet this phenomenon has not been fully explored across the STEM learning experience, where anxiety can impact negatively academic performance. Here, we evaluated anxiety and large-scale brain connectivity in 101 undergraduate physics students. We found sex differences in STEM-related and clinical anxiety, with longitudinal increases in science anxiety observed for both female and male students. Sex-specific relationships between STEM anxiety and brain connectivity emerged, with male students exhibiting distinct inter-network connectivity for STEM and clinical anxiety and female students demonstrating no significant within-sex correlations. Anxiety was negatively correlated with academic performance in sex-specific ways at both pre- and post-instruction. Moreover, math anxiety in male students mediated the relation between default mode-salience connectivity and course grade. Together, these results reveal complex sex differences in the neural mechanisms driving how anxiety is related to STEM learning.

---

Today’s universities and colleges are tasked with the challenge of developing novel strategies for improving undergraduate academic performance and ensuring that students are prepared for successful careers. In particular, emphasis is placed on enhancing student outcomes and generating enthusiasm for the science, technology, engineering, and mathematics (STEM) disciplines. However, STEM students encounter multiple, major-specific challenges, including intensive laboratory, project-based, and lecture-based coursework (Thiry et al., 2011), heightened classroom competition (Strenta et al., 1994; Gasiewski et al., 2012), and academic challenges of STEM courses (Strenta et al., 1994; Rask, 2010). As such, many students often struggle with STEM-related anxiety, which manifests as an unease, avoidance, or fear of learning science or math topics. In particular, female STEM students, relative to their male counterparts, are disproportionately affected by higher rates of STEM anxiety (Mallow, 1994; Brownlow et al., 2000; Baloglu and Kocak, 2006; Mallow et al., 2010). This may be due to STEM-related barriers that adversely impact achievement and performance (Kiefer and Sekaquaptewa, 2007; Nosek et al., 2009), including stereotype threat (Shapiro and Williams, 2012), gender-based bias (Moss-Racusin et al., 2012), and lack of non-stereotypical role models (Cheryan et al., 2011; Hernandez et al., 2018).

Despite the wealth of literature regarding STEM anxiety, little work has characterized the large-scale brain networks that may be linked with this barrier to learning and achievement in STEM students. However, significant prior neuroimaging research has contributed to our understanding of the neurobiological substrates of clinical anxiety and related psychiatric disorders (for reviews see: e.g., Peterson et al., 2014; Mochcovitch et al., 2014; Williams et al., 2017; Kim et al., 2018). In the context of psychopathology, a relatively recent paradigm shift from functional localization studies to large-scale brain network studies has occurred. Psychopathological processes, especially those found in mood disorders, are associated with aberrant organization and functioning of three key networks. First, the salience network (SN), anchored in the dorsal anterior cingulate cortex and frontoinsular cortex, plays a critical role in saliency detection, and attentional capture (Seeley et al., 2007; Menon and Uddin, 2010). Second, the default mode network (DMN), which includes the major nodes of the posterior cingulate and medial prefrontal cortices, is involved in self-referential processes and typically deactivates during stimulus-driven cognitive tasks (Grecius et al., 2003; Raichle, 2015). Third, the central executive network (CEN) is a frontoparietal system that includes the dorsolateral prefrontal and lateral posterior parietal cortices and is involved with cognitive processes such as working memory, problem solving, and goal-directed behavior (Dosenbach et al., 2007; Seeley et al., 2007). The interactions of these three large-scale networks underlies a unifying tripartite network model that seeks to characterize the maladaptive network organization and function common across psychiatric disorders (Menon, 2011; Sha et a., 2018). Within anxiety-related disorders, increased interactions between the SN, DMN, and CEN have been consistently observed (Sripada et al., 2012; Zhang et al., 2015) and SN-CEN and DMN-SN disruptions have been associated with trait anxiety in obsessive compulsive disorder (Fan et al., 2017) and diagnostic status in social anxiety disorder (Rabany et al., 2017). As hallmarks of STEM anxiety are similar to those of clinical anxiety (i.e., rumination, avoidance, over-generalization of threat stimuli), we expect these same large-scale networks to underlie anxiety in STEM students.

Here, we sought to bridge these research domains by examining the neurobiological correlates of STEM anxiety using the tripartite network model and its noted dysfunction in the context of clinical anxiety as a starting point. Given prior evidence in sex differences in STEM anxiety (Mallow, 1994; Brownlow et al., 2000; Baloglu and Kocak, 2006; Mallow et al., 2010), the present study investigated their neural substrates to advance towards a more complete model of anxiety-related mechanisms and strategies associated with learning processes. We examined if functional connectivity between the SN, DMN, and CEN is associated with STEM anxiety and whether this may differ among female and male STEM students. To this end, we collected self-report questionnaire and neuroimaging data from 101 university students (46F, 55M) who enrolled in and completed the first semester of a two-semester sequence of calculus-based, introductory physics. Introductory physics is a core “gateway” course on Newtonian mechanics and is required for undergraduate students seeking a university degree across a broad range of STEM fields, including chemistry, physics, engineering, or mathematics. Students completed behavioral and resting state functional magnetic resonance imaging (rs-fMRI) sessions at the beginning (pre-instruction) and ending (post-instruction) of the course. A robust body of evidence indicates that visuospatial ability (Pallrand and Seeber,1984; Kozhevnikov et al., 2002; Kozhevnikov and Thornton, 2006; Kozhevnikov et al., 2007) and mathematical competency (Cohen et al., 1978; Basson, 2002; Hudson and Liberman, 2005; Dehipawala et al., 2014; Korpershoek et al., 2015) are associated with and may predict physics learning and academic performance. Since science, spatial, and math anxiety may impede performance (Hembree, 1990; Vitasari et al., 2010; Núñez-Peña et al., 2013), we administered questionnaires probing science anxiety (Mallow, 1994), spatial anxiety (Lawton, 1994), and math anxiety (Alexander and Matray, 1989) collectively assess STEM-related anxiety. In addition, the Beck anxiety inventory was completed to assess clinical anxiety symptoms (Beck et al., 1988). To examine the relationships among STEM anxiety, brain connectivity, and sex, we addressed the following fundamental questions. First, are there sex differences in anxiety scores? Second, is there a relationship between STEM and clinical anxiety and functional connectivity? Third, are anxiety scores correlated with academic performance? Finally, does anxiety mediate the relationship between functional connectivity and academic performance? We predicted that anxiety scores would be significantly higher for female versus male STEM students. We also anticipated that functional connectivity would be correlated with STEM anxiety among both females and males, particularly when considering the SN. Finally, we hypothesized that STEM anxiety would be negatively correlated with academic performance for both female and male STEM students.

## RESULTS

### Sex differences in STEM anxiety

We performed mixed model ANOVA analyses for each anxiety measure^1^. These analyses demonstrated significant main effects of sex on all measures of anxiety, including science, spatial, math, and clinical anxiety (**Table 1**). Female students reported higher mean levels of anxiety on every measure compared to male students at both pre- and post-instruction (**Fig. 1**). When considering how students’ anxiety changed across the semester-long course, only science anxiety displayed a main effect of time. Examining the marginal means for female students, science anxiety scores were significantly increased at post-instruction (M = 16.43, SD = 10.76) compared to pre-instruction (M = 6.41, SD = 7.96). Similar results were observed for male students: science anxiety scores were significantly increased at post-instruction (M = 11.28, SD = 9.563) compared to pre-instruction (M = 3.15, SD = 3.498). There was no significant interaction between participant sex and change in anxiety scores on any measure.

**Table 1.** Results of Between-by-Within ANOVA on Anxiety Measures.

| Factor | Science |  |  | Spatial |  |  | Math |  |  | Clinical |  |  |
| --- | --- | --- | --- | --- | --- | --- | --- | --- | --- | --- | --- | --- |
| | $F$ | $p$ | $\eta^2_{\text{Partial}}$ | $F$ | $p$ | $\eta^2_{\text{Partial}}$ | $F$ | $p$ | $\eta^2_{\text{Partial}}$ | $F$ | $p$ | $\eta^2_{\text{Partial}}$ |
| Sex | 9.08 | 0.003 | 0.08 | 9.48 | 0.003 | 0.09 | 12.42 | 0.001 | 0.11 | 5.45 | 0.022 | 0.05 |
| Time | 101.52 | < 0.001 | 0.51 | 0.09 | 0.763 | 0.00 | 0.38 | 0.538 | 0.00 | 0.04 | 0.848 | 0.00 |

**Table 1.** Results of Between-by-Within ANOVA on Anxiety Measures.
|  |  |  |  |  |  |  |  |  |  |  |  |  |
| --- | --- | --- | --- | --- | --- | --- | --- | --- | --- | --- | --- | --- |
| Interaction | 1.10 | 0.297 | 0.01 | 0.76 | 0.387 | 0.01 | 0.20 | 0.657 | 0.00 | 1.17 | 0.282 | 0.01 |

Note: $\eta^2_{\text{partial}}$ is reported as calculated by SPSS. Numerator and denominator degrees of freedom for all $F$ ratios were 1 and 99, respectively.

**Fig. 1.**
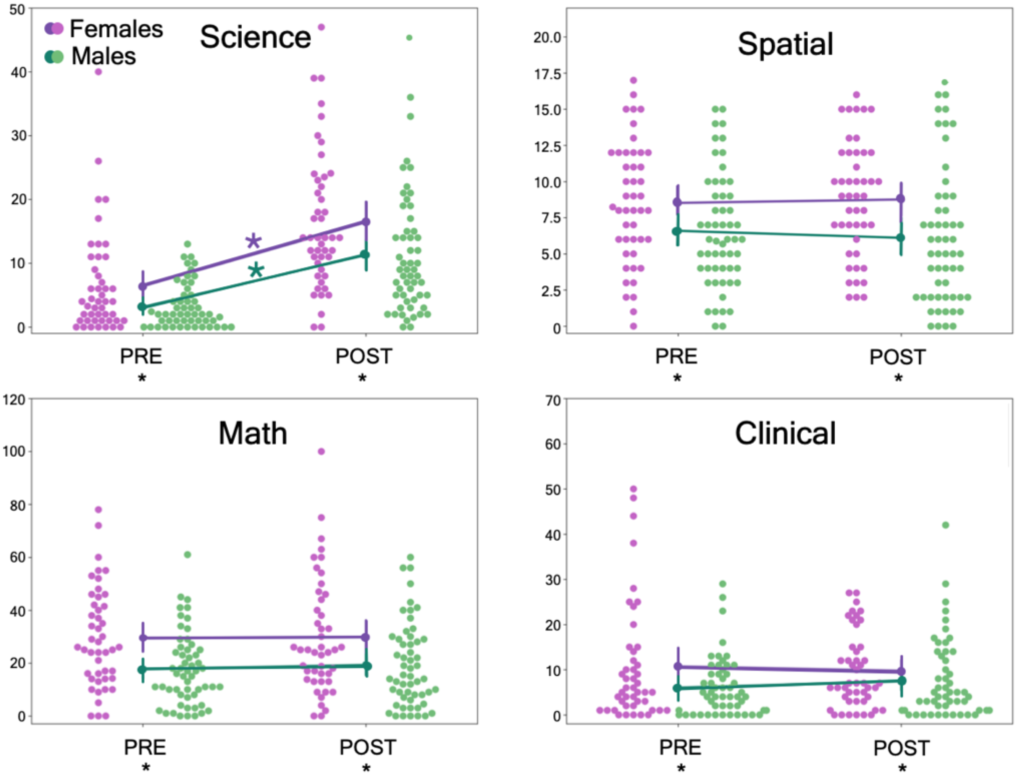
Sex Differences in Anxiety. Raw scores for science, spatial, math, and clinical anxiety (as measured by the Beck anxiety inventory) for female (purple) and male (green) undergraduate students enrolled in an introductory physics course. Anxiety was assessed at the beginning of the semester (i.e., pre-instruction or “PRE”) and at the completion of the course (i.e., post-instruction or “POST”). Black asterisks on bottom PRE/POST labels indicate significant sex differences in anxiety at PRE or POST. Purple and green asterisks indicate significant increases in science anxiety across time.

### Neural correlates of anxiety

To assess how functional brain connectivity relates to anxiety, we first identified the SN, DMN, and CEN using a data-driven, meta-analytic parcellation (Laird et al, 2011) (**Fig. 2**), extracted the average network time series from pre-processed rs-fMRI data, and constructed per-participant adjacency matrices reflecting the degree of between-network correlation across the three networks (Abraham et al., 2014). Motion was regressed out and high-motion volumes were censored (Power et al., 2014). The edge weights between the tripartite network connections were calculated as Pearson’s correlation coefficients between each network time series (e.g., inter-network functional connectivity between CEN-DMN, DMN-SN, and SN-CEN).

**Fig. 2.**
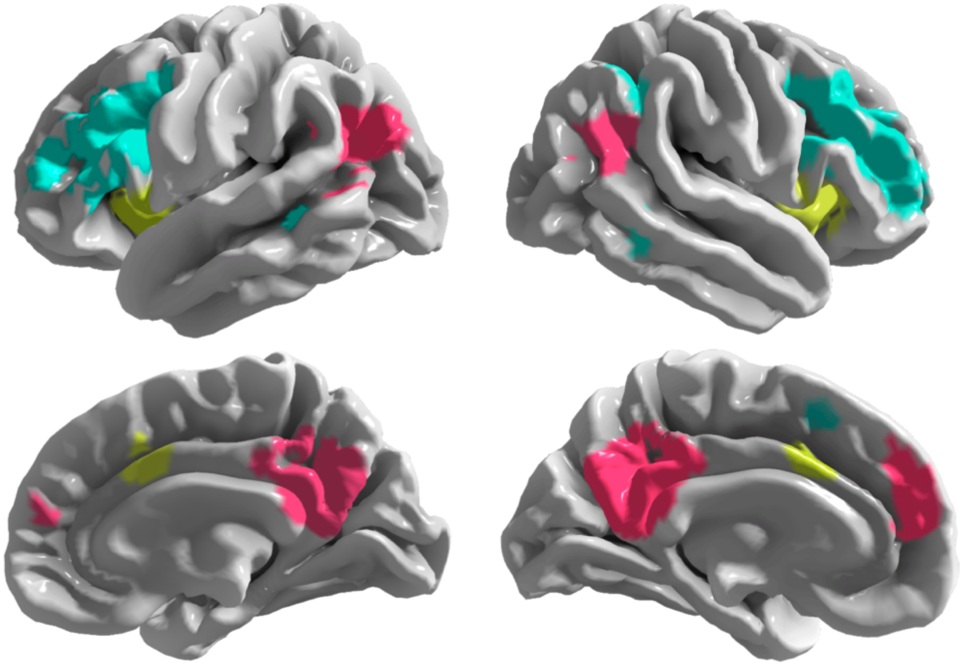
Network Parcellation. Network masks for the central executive (cyan), default mode (pink), and salience (yellow) networks were adapted from a data-driven, meta-analytic parcellation^41^ and used to extra network-wise signals from pre-processed rs-fMRI data from each participant.

To quantify putative relations between functional connectivity and anxiety, Pearson correlation coefficients were computed between the inter-network edge weights and anxiety scores, controlling for a false discovery rate of 0.25 using the Benjamini-Hochberg Procedure (Benjamini and Hochberg, 1995) (**Fig. 3**). At pre-instruction, among female students, there were no significant correlations between any of the anxiety scores and inter-network connectivity. In contrast, male students at pre-instruction exhibited significant correlations between science anxiety and CEN-DMN connectivity (*r*(53) = 0.275, *P* = 0.042, *α*_*FDR*_ = 0.13), science anxiety and DMN-SN (*r* = 0.311, *P* = 0.021, *α*_*FDR*_ = 0.10), spatial anxiety and CEN-DMN (*r* = 0.366, *P* = 0.006, *α*_*FDR*_ = 0.02), math anxiety and CEN-DMN (*r* = 0.325, *P* = 0.015, *α*_*FDR*_ = 0.08), math anxiety and DMN-SN (*r* = 0.355, *P* = 0.008, *α*_*FDR*_ = 0.04), and clinical anxiety and SN-CEN (*r* = −0.343, *P* = 0.010, *α*_*FDR*_ = 0.06). The correlation between clinical anxiety and SN-CEN connectivity was the only significant negative correlation observed, as well as the only measure linked with SN-CEN connectivity. All STEM anxiety measures in males were positively correlated with the CEN-DMN and DMN-SN connectivity. We also tested for an effect of sex across these results and observed that the correlation between clinical anxiety and SN-CEN was significantly different between female and male students (*Z* = −2.927, *P* = 0.002).

**Fig. 3.**
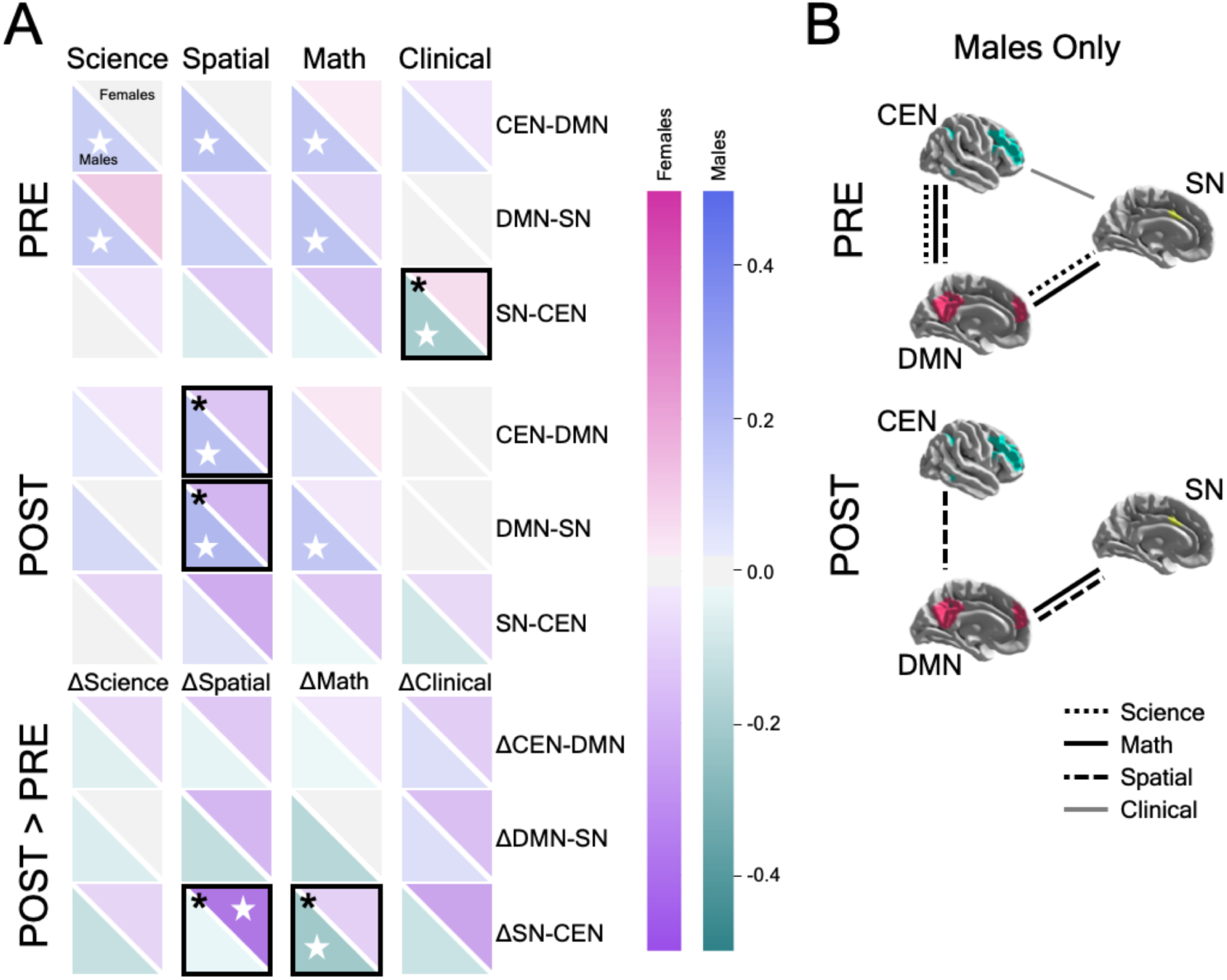
Anxiety and Functional Brain Connectivity. (A) Correlation values are shown between science, spatial, math, and clinical anxiety (columns) and between-network tripartite connectivity between the SN, DMN, and CEN networks (rows). Correlations are displayed for pre-instruction (“PRE”), post-instruction (“POST”), and the change across time (“POST > PRE”). Each square represents the correlation between anxiety and inter-network connectivity, with the upper diagonal displaying the value for female students and the lower diagonal representing male students. Positive and negative correlations are indicated by the color bars. Significant within-sex correlations are indicated by a white star, while significant between-sex correlations are indicated by a black box with an asterisk. (B) An alternative visualization of the results is provided to delineate the between-network correlations with anxiety in male students. While female students exhibited no significant correlations between anxiety and brain connectivity at pre- or post-instruction, male students exhibited several significant correlations at both time points. Males exhibited a general tendency to show fewer significant correlations at post-compared to pre-instruction associated with a reduced set of tripartite connections.

At post-instruction, no significant correlations were observed between anxiety scores and inter-network connectivity for female students. Male students at post-instruction exhibited significant correlations between spatial anxiety and CEN-DMN (r(53) = 0.381, P = 0.004, *α*_*FDR*_ = 0.04), spatial anxiety and DMN-SN (r = 0.435, P = 0.001, *α*_*FDR*_ = 0.02), and math anxiety and DMN-SN (r = 0.332, P = 0.013, *α*_*FDR*_ = 0.06). As with pre-instruction results, the significant STEM-related correlations were positive and only significantly related to the CEN-DMN and DMN-SN, but not SN-CEN connectivity. Again, we also tested for an effect of sex across these results and observed that the spatial anxiety correlations with CEN-DMN and DMN-SN and significantly differed between female and male students (Z = −2.375, P = 0.009 and Z = 3.094, P = 0.001, respectively).

In addition, we examined the correlations between the change in anxiety scores and the change in connectivity from pre- to post-instruction (detailed scatterplots shown in **Fig. S2**). Of these, Δanxiety_spatial_ and ΔSN-CEN were significantly negatively correlated for females (*r*(44) = −0.459, *P* = 0.001, *α*_*FDR*_ = 0.02), but not males *r*(53) = −0.041, *P* = 0.764, *α*_*FDR*_ = 0.23), and the difference between sexes was statistically significant, *Z* = 2.208, *P* = 0.014. Thus, for female students, as spatial anxiety increased over time, connectivity between SN and CEN decreased. Conversely, Δanxiety_math_ and ΔSN-CEN were significantly negatively correlated among male students (*r*(53) = −0.361, *P* = 0.007, *α*_*FDR*_ = 0.02), but not female students *r*(44) = −0.057, *P* = 0.707, *α*_*FDR*_ = 0.17), and this difference between sexes was statistically significant, *Z* = −1.557, *P* = 0.06. Thus, for male students, as math anxiety increased over time, connectivity between the SN and CEN decreased.

### Sex, anxiety, and academic performance

Traditional measures of academic performance include measures of students’ grades. We collected each student’s overall GPA prior to taking the course, as well as their final physics course grade. First year students were excluded (2F, 6M) from the GPA analysis since they entered the physics course with a GPA of zero. No significant sex differences were observed for incoming GPA (*U*_*GPA*_ = 1051.5, *P* = 0.838, *d* = 0.293) or physics course grade (*U*_*grade*_ = 1056.5, *P* = 0.148, *d* = 0.286).

To quantify the relation between anxiety and academic performance, Pearson correlations were computed separately for female and male students, controlling for a false discovery rate of 0.25 using the Benjamini-Hochberg Procedure (Benjamini and Hochberg, 1995) (**Fig. 4**). Among female students at pre-instruction, GPA was positively correlated with spatial anxiety (*r*(42) = 0.381, *P* = 0.011, *α*_*FDR*_ = 0.06) while course grade was negatively correlated with math anxiety (*r*(44) = −0.321, *P* = 0.030, *α*_*FDR*_ = 0.09) and clinical anxiety (*r*(44) = −0.534, *P* < 0.001, *α*_*FDR*_ = 0.03). Among male students at pre-instruction, GPA was only negatively correlated with math anxiety (*r*(47) = −0.358, *P* = 0.012, *α*_*FDR*_ = 0.03). The correlation between GPA and clinical anxiety at pre-instruction significantly differed between females and males (*Z* = 2.364, *P* = 0.009). Among female students at post-instruction, GPA was negatively correlated with clinical anxiety (*r*(42) = −0.315, *P* = 0.037, *α*_*FDR*_ = 0.06), and grade was negatively correlated with both math anxiety (*r*(44) = −0.293, *P* = 0.048, *α*_*FDR*_ = 0.09) and clinical anxiety (*r*(44) = −0.401, *P* = 0.006, *α*_*FDR*_ = 0.03). Among male students at post-instruction, GPA was negatively correlated with science anxiety (*r*(47) = −0.370, *P* = 0.009, *α*_*FDR*_ = 0.09) and math anxiety (*r*(47) = −0.449, *P* = 0.001, *α*_*FDR*_ = 0.03), and similarly, grade was also negatively correlated with science anxiety (*r*(53) = −0.354, *P* = 0.008, *α*_*FDR*_ = 0.06) and math anxiety (*r*(53) = −0.422, *P* = 0.001, *α*_*FDR*_ = 0.03). Thus, in general, high levels of post-instruction STEM anxiety were associated with poor academic performance. No significant sex differences at post-instruction were observed.

**Fig. 4.**
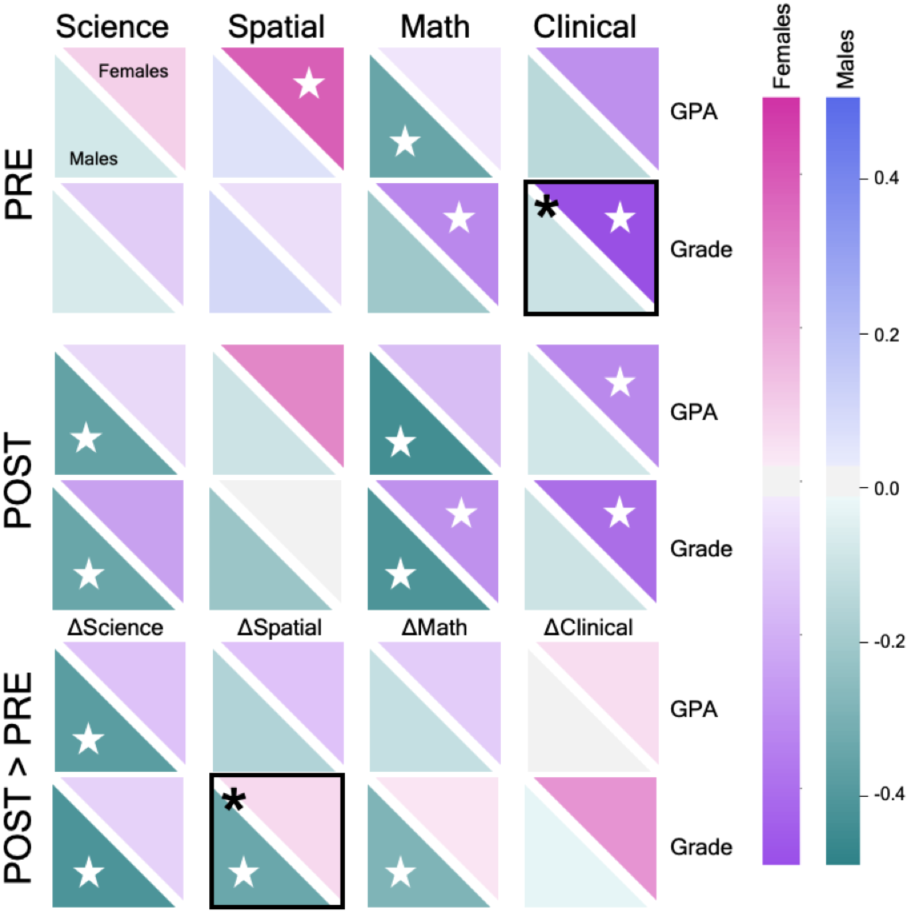
Sex, Anxiety, and Performance. Correlation values are shown between science, spatial, math, and clinical anxiety (columns) and pre-semester GPA and physics course grade (rows). Correlations are provided for pre-instruction (“PRE”), post-instruction (“POST”), and the change across time (“POST > PRE”). Each square represents the correlation between anxiety and GPA/grade, with the upper diagonal displaying the value for female students and the lower diagonal representing the male students. Positive and negative correlations are indicated by the color bars. Significant within-sex correlations are indicated by a white star, while significant between-sex correlations are indicated by a black box with an asterisk.

Next, we examined the correlations between the change in anxiety scores and academic performance. Female students demonstrated no significant correlations between GPA or grade and the change in any anxiety measure. Conversely, male students exhibited significant negative correlations between grade and Δanxiety_science_ (*r*(53) = −0.393, *P* = 0.003, *α*_*FDR*_ = 0.03), Δanxiety_spatial_ (*r* = −0.339, *P* = 0.011, *α*_*FDR*_ = 0.06), and Δanxiety_math_ (*r* = −0.296, *P* = 0.028, *α*_*FDR*_ = 0.09), as well as between GPA and Δanxiety_science_ (*r*(47) = −0.416, *P* = 0.003, *α*_*FDR*_ = 0.03). A significant sex effect was observed for the correlation between grade and Δanxiety_spatial_ (*Z* = −2.033, *P* = 0.021).

### Anxiety mediates brain function and performance

Lastly, we investigated if functional brain connectivity was correlated with academic performance at pre- or post-instruction, controlling for a false discovery rate of 0.25 using the Benjamini-Hochberg Procedure (Benjamini and Hochberg, 1995) (**Fig. 5a**). For female students, no significant correlations were observed between inter-network brain correlations and GPA or course grade at either time point. For male students, there was a significant, negative correlation between DMN-SN connectivity and course grade at post-instruction (*r*(53) = −0.267, *P* = 0.049, *α*_*FDR*_ = 0.09). Given this result, we then asked to what extent anxiety might mediate the relationship between brain connectivity and academic performance. We investigated four separate mediation models among male students to determine if post-instruction science, spatial, math, or clinical anxiety was a mediating variable on DMN-SN connectivity and course grade. We observed including math anxiety as a variable reduced the total effect of DMN-SN and course grade, which was no longer significant (*indirect effect* = - 0.544, *SE* = 0.267, *P* = 0.042; *95% bootstrap confidence intervals* (*CIs*) = −1.161, −0.128) (**Fig. 5b**). Science, spatial, and clinical anxiety were not found to mediate DMN-SN connectivity and course grade.

**Fig. 5.**
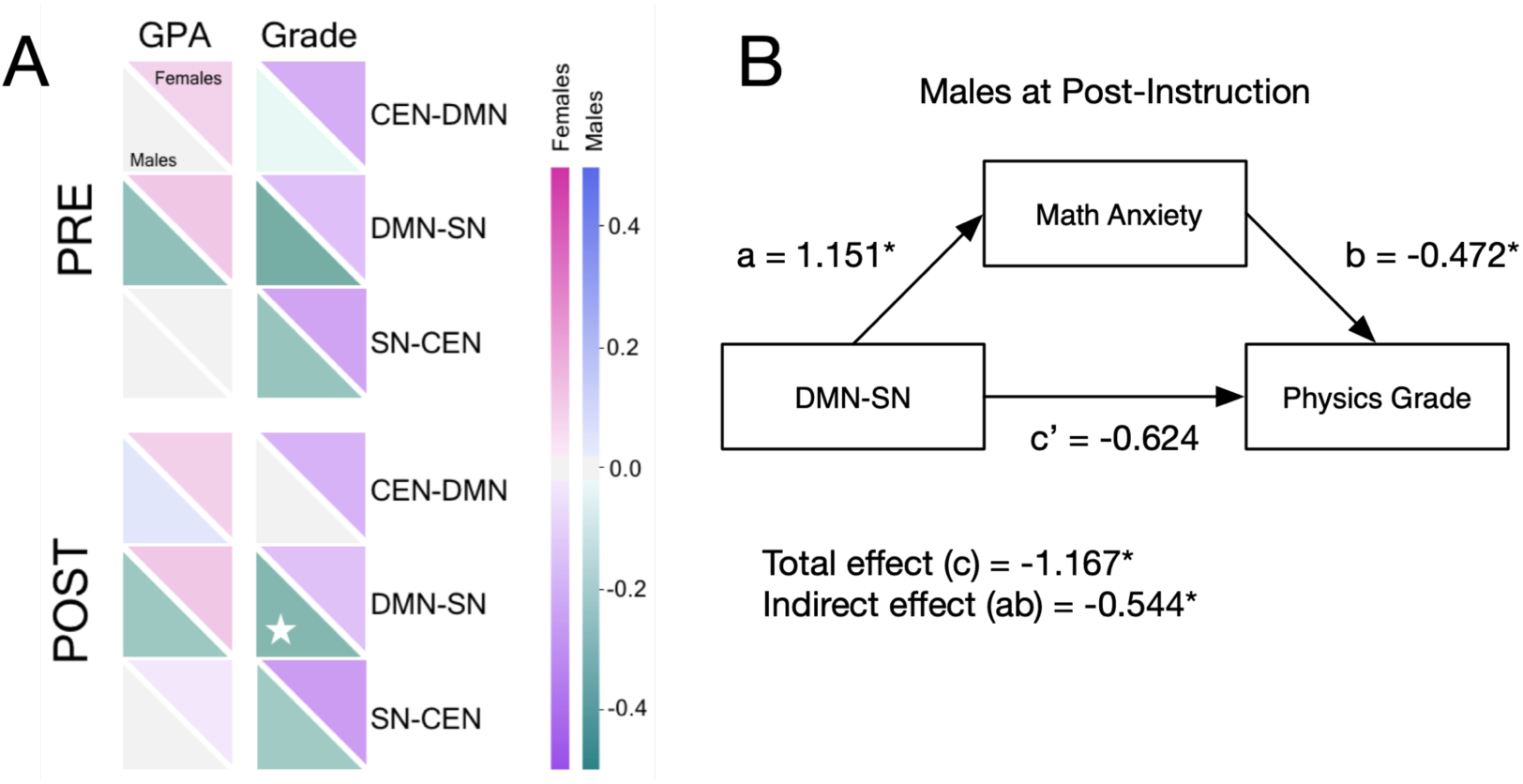
Post-instruction math anxiety mediates the relation between DMN-SN connectivity and physics course grade. (A) Correlation values are shown between pre-semester GPA and physics course grade (columns) and between-network tripartite connectivity between the SN, DMN, and CEN networks (rows). Correlations are provided for pre-instruction (“PRE”) and post-instruction (“POST”). Each square represents the correlation between GPA/grade and inter-network connectivity, with the upper diagonal displaying the value for female students and the lower diagonal representing the male students. Positive and negative correlations are indicated by the color bars. Significant within-sex correlations are indicated by a white star (B) Results of the mediation analysis indicated that every 1-unit increase in post-instruction DMN-SN connectivity was associated with a a = 1.151 (SE = 0.427, P = 0.007) unit increase in post-instruction math anxiety. Adjusting for post-instruction DMN-SN connectivity, every unit increase in post-instruction math anxiety was associated with a b = −0.472 (SE = 0.144, P = 0.001) unit decrease in course grade. Increases in post-instruction DMN-SN connectivity were associated with decreases in course grade, indirectly through increases in post-instruction math anxiety. Specifically, for every a = 1.151-unit increase in post-instruction math anxiety, there was a ab = −0.544 (SE = 0.267, P = 0.042) unit decrease in course grade. Importantly, a bias-corrected bootstrapped confidence interval with 10,000 samples (Rosseel et al., 2012) did not contain 0, 95% CI [−1.161, −0.128], indicating a significant indirect effect (ab). Last, there was no sufficient evidence that post-instruction DMN-SN connectivity was significantly associated with course grade, independent of its association with post-instruction math anxiety, c’ = - 0.624 (SE = 0.624, P = 0.318).

## DISCUSSION

Our results identified significant sex differences in STEM and clinical anxiety, among undergraduate physics students, with females experiencing higher levels of STEM anxiety compared to their male counterparts, in agreement with prior work (Alexander and Matray, 1989; Mallow, 1994; Lawton, 1994). While we observed significantly increased science anxiety from pre- to post-instruction in both female and male students, we found no evidence of an interaction between sex and change in anxiety scores. That is, our results do not suggest that the introductory physics course in our study differentially impacts changes in anxiety for female and male students. This is important from the perspective of educators who seek to create inclusive classrooms that are free from instructionally derived bias.

Previous studies have shown that SN, DMN, and CEN dysfunction are implicated in clinical anxiety (Sripada et al., 2012; Zhang et al., 2015; Fan et al., 2017). We were surprised to see that female students exhibited no significant correlations between connectivity and anxiety at either time point. In contrast, male students exhibited multiple, significant positive correlations between connectivity and STEM anxiety at both pre- and post-instruction and a negative correlation between clinical anxiety and SN-CEN at pre-instruction. Dynamic interactions between the SN, DMN, and CEN are critical for successful execution of a wide range of cognitive and emotional processes. Healthy inter-network equilibrium is thought to rely on suppression of self-referential cognition in the DMN (Gusnard et al., 2001) to allow for identification of salient, task-relevant stimuli in the SN that should be relayed to the CEN (Sridharan et al., 2008), resulting in anti-correlations between the DMN and CEN (Fox et al., 2005). Evidence suggests that increased anxiety is associated with *increased* functional connectivity between the SN and DMN in clinical anxiety disorders (Sripada et al., 2012; Zhang et al., 2015; Fan et al., 2017). In contrast, the converse relationship has also been observed: higher levels of trait anxiety in healthy adolescents are related to *decreased* functional connectivity of the SN to DMN and CEN regions (Geng et al., 2016). Our current results in male students suggest anxiety-related disruption of inter-network equilibrium between the SN, DMN, and CEN and provide additional STEM-relevant support for the importance of suppressing self-referential DMN interactions to maintain a healthy balance across networks. DMN-SN connectivity was negatively correlated with course grade in male students at post-instruction, further supporting the importance of toggling off internal processing when salient events are detected in the context of STEM learning.

Male students exhibited a general trend of fewer significant brain-anxiety correlations at post-compared to pre-instruction, despite increased science anxiety. Although speculative, this tendency is suggestive of a cognitive or physiological mechanism at play and may provide directions for future work. As male students are faced with the challenges of their first university-level physics course, the brain may accommodate the increases in science anxiety and balance the response to such challenges. In contrast, female students experience greater obstacles in STEM education that can trigger anxiety as early as the preschool and elementary years (Gunderson et al., 2012; Hill et al., 2016; Wong et al., 2017). The null female results may point to a lack of vulnerability, suggesting that their relatively higher STEM anxiety does not hinder salience-related central executive and self-referential processes. Female students may experience an earlier adaptive period as their STEM anxiety increases, resulting in a compensatory mechanism that down-regulates the anxiety-brain correlations, possibly via a reallocation of neural resources or a functional reorganization of anxiety-related systems. Overall, it is unclear if the sex differences in functional connectivity observed here reflect experiential differences in STEM anxiety-related developmental trajectories due to disruptions in emotion regulation (McRae et al., 2008), attentional control (Bishop et al., 2004; Gur et al., 2012; Roalf et al., 2014), motivation and drive (Freudenthaler et al., 2008; Bugler et al., 2015; Young et al., 2015), disengagement and avoidance (Panayoitou et al., 2017), coping strategy (Normann and Esborn, 2019) or a combination of these influences. Further work is needed to investigate sex differences in developmental STEM trajectories, to determine if female students experience STEM-related anxiety and learn strategies for counterbalancing their anxiety at an earlier educational stage.

Aberrant connectivity between the CEN and SN in anxious individuals may result from a diminished ability to exert cognitive control and regulate emotional responses (Menon and Uddin, 2010). Previous work has shown that university students with high math anxiety exhibit increased SN activity when anticipating a math problem (Lyons and Beilock, 2012), yet math cue-related activity increased in the CEN as math deficit decreased, suggesting that increased recruitment of cognitive control processes may improve performance in math (Lyons and Beilock, 2011). Relatedly, lower math anxious children showed increased activation in regions of the CEN and DMN during math problem solving compared to higher math anxious children (Young et al., 2012) although the reverse was shown by Supekar et al. (2015) during successful math trials. This prior work in task-based fMRI has not addressed sex-related differences in the neural correlates of anxiety. Here, we showed math anxiety was consistently related to brain connectivity and performance for both sexes compared to other anxiety measures. Specifically, although math anxiety was not significantly related to SN-CEN inter-network connectivity in male students at pre- or post-instruction, the change in math anxiety was negatively correlated with the change in SN-CEN connectivity over the course of instruction. That is, as math anxiety increased across the semester for male students, SN-CEN connectivity also increased. Although higher levels of math anxiety are reported by female students, math anxiety has been more strongly linked to poor performance in precollege male students (Hembree, 1990). Our results related to math anxiety in male students suggest that the SN-CEN pathway may play a critical role in longitudinal changes across a semester of STEM learning, but that the DMN-SN pathway is more strongly related to course performance, with math anxiety mediating this relationship.

Our study is limited by several concerns. First, our objective was to characterize sex differences in STEM anxiety in STEM undergraduate students. As such, recruitment and enrollment of participants who completed a core STEM course required broadly across STEM majors was deemed a key aspect of this study – our target sample was a wide range of STEM undergraduates, which we captured via an introductory physics course. However, it is likely that our results do not generalize to non-STEM undergraduates, given their different experiences with STEM-related coursework. Future work is needed to clarify how STEM anxiety may be differentially experienced by non-STEM students compared to STEM students. Second, students diagnosed with psychiatric or neurologic disorders were excluded; participants were also excluded if they reported use of psychotropic medications. Thus, our results may not generalize to a broader community of students that includes those diagnosed with and receiving treatment for clinical disorders of anxiety and depression. Third, although our primary analyses treated STEM and clinical anxiety as independent constructs, we acknowledge that this may not be the case for some students. We conducted collinearity diagnostics, which demonstrated that multicollinearity was not a concern for STEM and clinical anxiety measures. As an added step to reduce potential confounds by clinical anxiety, we performed partial Pearson correlation analyses that produced approximately equal, and even in some instances stronger, associations between STEM anxiety, functional connectivity, and academic performance when controlling for clinical anxiety. Both the collinearity diagnostics and the additional partial correlation analyses are available in the Supplemental Information (SI). Fourth, the timeline of the study created logistic challenges in that all data collection was carried out during short periods of time at the beginning and ending of each semester. As a result, while MRI sessions were completed following the final exam, our post-instruction behavioral data were generally scheduled the week prior to finals week (a period of time generally associated with increased anxiety levels among students). It is unclear how our results may be confounded by the temporal mismatch of MRI and behavioral sessions. Fifth, additional clarity may have been provided by including additional measures (e.g., the Positive and Negative Affect Schedule) to assess participant mood states on the day of scanning. Moreover, MRI scans may induce anxiety for some participants, especially those with high trait anxiety. Future work should strongly consider including measures of MRI-related anxiety (e.g., the Magnetic Resonance Imaging-Anxiety Questionnaire (Ahlander et al., 2016)). Last, anxiety was assessed exclusively via self-report rating scales. Future work should include additional multi-method designs such as task-based fMRI with concurrent psychophysiological indexes of sympathetic and parasympathetic activity (e.g., respiratory sinus arrhythmia and skin conductance, respectively).

Overall, our results indicate that female and male students experience different levels of STEM anxiety and exhibit different neurobiological systems-level support for this anxiety, which is differentially associated with their academic success. That this occurs despite no sex differences in performance (e.g., GPA or course grade) is notable, and in agreement with two recent meta-analyses that provide strong evidence challenging the persistent stereotypes that male students possess higher innate aptitude in math and science compared to female students (Kersey et al., 2018; O’Dea et al., 2018). Importantly, the course studied here was shown to be equal (i.e., no significant interaction between sex and change in anxiety), but not equitable (i.e., did not reduce sex differences). The gender gap in STEM remains largely unexplained (Riegle-Crumb et al., 2012), yet our results suggest that female students maintain performance compared to their male counterparts while responding differently to obstacles and challenges associated with STEM learning. Organizations supporting women in STEM have long promoted the idea that reduced female representation in STEM is due to poor climate for women rather than lack of ability or interest. Our results support this framework. We recommend that positive changes in favor of promoting women in STEM should focus on addressing climate issues that contribute to STEM anxiety. At the elementary and secondary school level this could include improving parental and teacher support, which has been shown to significantly impact girls’ anxiety, confidence, and performance (Beilock et al., 2010; Gunderson et al., 2012; Casad et al., 2015). At the university level, this could include increasing visible role models (e.g., women as STEM faculty and in senior leadership positions; Winslow and Davis, 2016), revising ineffective Title IX policies (a United States statute that protects students from sex-based discrimination in federally-funded education programs and activities; US Department of Education, 2015), and enacting a zero-tolerance policy for sexual harassment and abuse at institutions, research societies, and federal funding agencies. It is incumbent upon university leaders to optimize pathways for all students entering the national STEM workforce. Instructional techniques focused on helping students learn content while building positive affect may be of particular importance in supporting learning that is inclusive for all students, thereby retaining individuals that drop out of STEM careers due to these climate-related factors. Continued development of instructional practices should emphasize the important distinction between equality and equity.

Broadly, female and male STEM students experience different learning environments, societal expectations, and academic opportunities, which all contribute to socio-emotional brain development, necessitating rigorous and objective standards for the study of sex and gender in neuroimaging research (Rippon et al., 2014). Our results demonstrate that sex differences in brain networks are not fixed and that STEM anxiety is related to changes in both female and male students’ brains during the physics learning process. We conclude that there are significant sex differences between STEM anxiety linked with large-scale brain networks and recommend future research to determine how reducing barriers and making the climate more equitable may enable a more inclusive STEM community.

## METHODS

### Participants and Study Design

One hundred and one healthy right-handed undergraduate students (mean age = 19.94 ± 2.46 years, range = 18-25 years; 46 females) who completed a semester of introductory calculus-based physics at Florida International University (FIU) took part in this study. Participants self-reported that they were free from cognitive impairments, neurological and psychiatric conditions, and did not use psychotropic medications. The physics course emphasized problem solving skill development and covered topics in classical Newtonian mechanics, including motion along straight lines and in two and three dimensions, Newton’s laws of motion, work and energy, momentum and collisions, and rotational dynamics. Students completed a behavioral and MRI session at two time points at the beginning (“pre-instruction”) and conclusion (“post-instruction”) of the 15-week semester. Pre-instruction data collection sessions were generally acquired no later than the fourth week of classes. Post-instruction sessions were completed no more than two weeks after the final exam. Written informed consent was obtained in accordance with FIU’s Institutional Review Board approval.

### Behavioral Measures

Participants completed a series of self-report instruments during their pre- and post-instruction behavior session, including, but not limited to: the Science Anxiety Questionnaire (Mallow, 1994), the Spatial Anxiety Scale (Lawton, 1994), the Mathematics Anxiety Rating Scale (Alexander and Matray, 1989), and the Beck Anxiety Inventory (Beck et al., 1988). Tests were performed to determine if our data on science, spatial, math, and clinical anxiety met the assumption of collinearity and the results indicated that multicollinearity was not a concern; collinearity diagnostics are provided in the SI. Participants also provided their demographic details (e.g., biological sex, age).

### Missing Data

A missing value analysis indicated that less than 2% of the data were missing for each variable and these were observed to be missing completely at random (MCAR). We chose not to implement multiple imputation, expectation maximization, or regression because the data violated the assumption of multivariate normality (Dong and Peng, 2013). Given the small sample size, frequency of missingness (1-2%), and lack of systematic reasons for missingness, we implemented item-level mean substitution imputation to avoid case-wise deletion of missing data (Rubin et al., 2007).

### fMRI Acquisition and Pre-Processing

Neuroimaging data were acquired on a GE 3T Healthcare Discovery 750W MRI scanner at the University of Miami. Resting state functional MRI (rs-fMRI) data were acquired with an interleaved gradient-echo, echo planar imaging (EPI) sequence (TR/TE = 2000/30ms, flip angle = 75°, field of view (FOV) = 220×220mm, matrix size = 64×64, voxels dimensions = 3.4×3.4×3.4mm, 42 axial oblique slices). During resting-state scans participants were instructed to remain still with their eyes closed. A T1-weighted series was also acquired using a 3D fast spoiled gradient recall brain volume (FSPGR BRAVO) sequence with 186 contiguous sagittal slices (TI = 650ms, bandwidth = 25.0kHz, flip angle = 12°, FOV = 256×256mm, and slice thickness = 1.0mm). Each participant’s structural T1-weighted image was oriented to the MNI152 2mm template using AFNI’s (http://afni.nimh.nih.gov/afni; Cox, 1996) 3dresample, then skull-stripped using the Brain Extraction Tool from FMRIB’s Software Library (FSL, https://fsl.fmrib.ox.ac.uk/fsl/fslwiki; Smith et al., *2002*; Jenkinson et al., 2012). Utilizing FSL’s automated segmentation tool (FAST), tissue-type masks were generated to inform nuisance parameters (Zhang et al., 2001). Then, utilizing FSL’s FLIRT (Jenkinson and Smith, 2001), the middle volume of each functional run was extracted and coregistered with the corresponding T1-weighted image. Utilizing FSL’s MCFLIRT with spline interpolation, motion correction aligned all volumes of each subject’s rs-fMRI time series with that middle volume. To further correct for in-scanner motion effects, functional volumes unduly affected by motion were identified using fsl_motion_outliers, with a framewise displacement threshold of 0.2mm (Power et al., 2014). Resultant motion artifacts were removed with ICA-AROMA (https://github.com/rhr-pruim/ICA-AROMA; Pruim et al., 2015). Then, CSF and WM masks were transformed into functional native space, eroded by 1 and 2 voxels, respectively, and from each the mean signal was extracted and used to regress out non-neural signals in a final nuisance regression step using AFNI’s 3dTproject, which detrended and normalized the rs-fMRI time series, as well. Finally, rs-fMRI images were transformed into MNI152 2mm space for further data analysis.

### Network Parcellation and Brain Connectivity Analyses

Each participant’s rs-fMRI data were standardized and parcellated according to the meta-analytic network components described by Laird et al. (2011). Included in this parcellation are the salience network (SN), default mode network (DMN), and central executive network (CEN). As these networks were delineated via ICA, some overlap was present between component maps. This overlap was resolved by a combination of proportional thresholding and manual editing, performed with the Mango image analysis tool (v. 4.0.1, http://ric.uthscsa.edu/mango/); final networks are shown in **Fig. 2**. Adjacency matrices were constructed per participant using Nilearn (v. 0.3.1, http://nilearn.github.io/index.html), a Python (v 2.7.13) module, built on scikit-learn, for the statistical analysis of neuroimaging data (Abraham et al., 2014; Pedregosa et al., 2011). For each of the three networks of interest, a single time series was computed as an average of the rs-fMRI time series from all voxels within the network, after further regressing out six motion parameters (from MCFLIRT) and censoring high-motion volumes (framewise displacement >0.2mm), as well as the immediately preceding volume and two following volumes, following recommendations from Power et al. (2014). Edge weights for each graph were Pearson’s correlations, calculated pairwise for the three networks, which are the graph’s nodes, resulting in a 3×3 network-wise correlation matrix for each participant. Although our emphasis focused on characterizing the putative relationships between inter-network connectivity and anxiety, we additionally analyzed intra-network connectivity to explore the relationship between within-network cohesion and anxiety. Pairwise correlation coefficients between constituent nodes of the SN, DMN, and CEN were computed and averaged within each network to obtain measures of intra-network cohesion. Pearson correlation coefficients were calculated between intra-network cohesion and anxiety scores, including science, spatial, math, and clinical anxiety. Among both female and male students, no significant relationships were observed between intra-network cohesion and anxiety within the SN, DMN, or CEN at either pre- or post-instruction.

### Statistical Analyses

All statistical tests were computed using IBM SPSS software, R Statistical Software, and Python tools/packages including Nilearn: Machine learning for Neuroimaging in Python, pandas (Python Data Analysis Library), matplotlib, Seaborn: statistical data visualization, Statsmodels, and SciPy. Observed *P* values are reported for statistical comparisons deemed significant after controlling for a false discovery rate of 0.25 using the Benjamini-Hochberg Procedure (Benjamini and Hochberg, 1995). The choice of the family of inferences over which an error rate is controlled is often ambiguous and a topic of scholarly debate (Holland and Cheung, 2002). In our study, we applied the Benjamini-Hochberg correction to each specific research question and assumed independence for each group and time point. For example, for the question “*What brain connections (3) correlate with anxiety (4) at pre-instruction for female students*?”, we corrected for 12 tests. We utilized adjusted alpha levels for each family of comparisons to impose a more conservative criterion for significance and avoid Type I errors.

## Data Availability

A GitHub repository was created at http://github.com/nbclab/PhysicsLearning/tree/master/anxiety to archive the source files for this study, including data analysis processing scripts and behavioral data. The network masks for the bilateral SN, DMN, and CEN are available via NeuroVault at https://neurovault.org/collections/4727/.

## ACKNOWLEDGMENTS

Primary funding for this project was provided by NSF REAL DRL-1420627; additional support to various authors was provided by NSF 1631325, NIH R01 DA041353, NIH U01 DA041156, NSF CNS 1532061, NIH K01DA037819, NIH U54MD012393, and the FIU Graduate School Dissertation Year Fellowships. Thanks to Karina Falcone, Rosario Pintos Lobo, and Camila Uzcategui for their assistance with data collection and to the Department of Psychology of the University of Miami for providing access to their MRI scanner. Special thanks to the FIU undergraduate students who volunteered and participated in this project.

## AUTHOR CONTRIBUTIONS

ARL, EB, SMP, MTS, RWL conceived and designed the project. JEB, EIB, RO acquired behavioral and fMRI data. AG, KLB, JEB, ARL analyzed data. KLB, MCR, TS contributed scripts and pipelines. AG, KLB, JEB, ARL wrote the paper. ARL contributed to all aspects of the project.

## Footnotes

1 *To assess the robustness of these results to potential violations of the assumptions of normality and equal variances, we replicated the analyses above using robust ANOVA methods recommended by Wilcox (2017). The robust ANOVAs returned the same pattern of results as the classical ANOVAs, strengthening our confidence in these findings. Additionally, we note that the same general pattern of results held when running the analyses using linear mixed model (multilevel) regressions, both with and without controlling for clinical anxiety when analyzing the remaining anxiety measures, available in Table S2 of the Supplementary Information (SI).*

