## Supplementary Information for "Sex differences in brain correlates of STEM anxiety"

### SUPPLEMENTAL INFORMATION

**Participants.** Female and male students who participated in this study were enrolled in and completed the first semester of a two-semester sequence of calculus-based, introductory physics (PHY 2048) at Florida International University (Miami, FL). This is a "gateway" course on Newtonian mechanics and is required for undergraduate students seeking a university degree across a broad range of STEM fields, including chemistry, physics, engineering, or mathematics. Importantly, this is a required course for STEM majors not just at FIU, but at universities throughout the US and international community. Eligible participants had never completed a university-level physics course and met a minimum grade point average (GPA) of 2.25. A GPA threshold of 2.25 (on a 4.0 scale) was chosen to identify students who were likely to complete the course and participate in the study at both time points. Generally, grade point averages of 2.0 are considered to be 'average' and are equivalent to a 'C' letter grade.

Participants self-reported that they were free from cognitive impairments, had not received a medical diagnosis for a neurological or psychiatric condition, did not use psychotropic medications, and presented at the first scan with no MRI contraindications. 129 individuals were consented into the study. During the pre-instruction data acquisition period (i.e., behavioral and MRI), 11 students were removed from the study due to scheduling conflicts or ineligibility concerns, yielding pre-instruction data from 118 healthy, right-handed (0.5 or greater on the Edinburgh Handedness Inventory; Oldfield, 1971), English-speaking undergraduate students (mean age = 20.16 ± 2.32 years, range = 18-25 years; 65 females. Following pre-instruction data collection, 3 students dropped the physics course during the semester and were removed from the study, 2 were non-responsive for post-instruction scheduling, 2 were no longer interested in participating, 2 were removed due to acquired MRI contraindications or ineligibility, and 1 student was unable to schedule their final MRI visit due to scheduling conflicts. Additionally, due to equipment malfunction, the post-instruction behavioral data from 4 participants and the post-instruction MRI data from 1 participant were lost due to data corruption. A total of 103 participants completed the study at post-instruction. Two data sets were removed from the analyses due to technical issues, yielding a final sample of 101 participants in the present study.

**Fig. S1** describes the missing study data across the stages of consent, data acquisition, and analysis. The final sample yielded matched (pre- and post-instruction) fMRI data sets from 101 students (mean age = 19.93 ± 2.46 years, range = 18-25 years; 46 females).

In line with scholarship on race and ethnicity (Phinney, 1996; Betancourt, 1993) and recommendations from the U.S. Census Bureau (2017a; 2017b), participants reported their racial self-identification with one or more groups, including, American Indian or Alaska Native, Asian, Black or African American, Native Hawaiian or Other Pacific Islander, White, and/or Other. The racial demographics of the sample were 3% American Indian or Alaskan Native, 9.9% Asian, 13.9% Black or African American, 0% Native Hawaiian or Other Pacific Islander, 72.3% White, and 6.9% Other. Even though there are various ethnic groups in the United States, in line with the U.S. Census Bureau, we used the question "*Are you of Spanish/Hispanic/Latinx ethnicity?*" to determine Spanish/Hispanic/Latinx identification. The racial demographics of the 65 participants (64 % of the total sample) who endorsed a Spanish/Hispanic/Latinx ethnicity were: 3% American Indian or Alaskan Native, 2.7% Asian, 2.7% Black or African American, 0% Native Hawaiian or Other Pacific Islander, 87.7% White, and 5.5% Other.

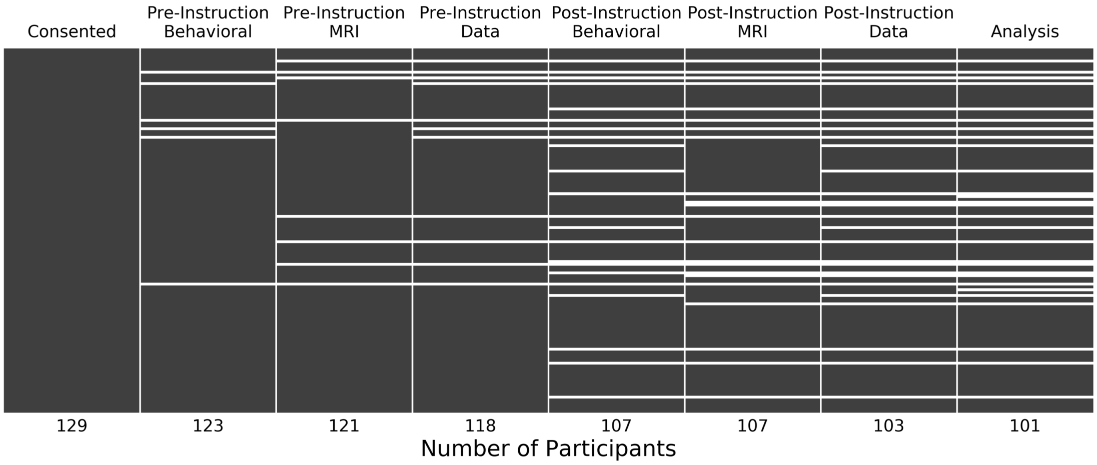


**Fig. S1.** Study Completeness Summary. *Display of the data table's nullity matrix for the study, describing missing data from study consent to all stages of pre- and post-instruction data collection and analysis. Visualization provided by* missingno*, a Python package for visualizing missing data (Bilogur, 2018).*

Behavioral data were acquired during the same time period with the exception of two individuals who completed their pre-instruction behavioral sessions in the fifth week of the semester. In general, post-instruction behavioral data were acquired two weeks prior to the start of final exams. However, ten students were unable to complete behavioral sessions within this timeline and instead completed their sessions during or after finals week. Post-instruction imaging sessions were held after the final exam of each physics course. In most cases this meant that students underwent MRI scanning in the two weeks following the University's final exam week. However, in some cases (e.g., due to conflicts in individual's post-semester travel schedules) students were unable to complete post-instruction MRI scanning after the semester ended. Thus, a total of 15 individuals completed post-instruction MRI scanning during finals week. For these sessions the study team attempted to schedule the MRI scan after the completion of the student's full set of finals exams, resulting in 13 students who completing their post-instruction MRI scan on the last day of finals week, after all exams had concluded.

**High Clinical Anxiety in Female Participants.** As shown in **Fig. 1**, there were four female participants with noticeably higher clinical anxiety at pre-instruction compared to the other participants. These four participants endorsed scores consistent with "potentially concerning levels of anxiety" (Beck et al., 1988), even though students diagnosed with psychiatric or neurologic disorders were excluded from the study. It is possible that some participants were experiencing clinical levels of anxiety that were undiagnosed prior to study enrollment. However, at post-instruction, these participants endorsed much lower anxiety scores, consistent with moderate levels of anxiety (Beck et al., 1988). Unfortunately, high levels of clinical anxiety in college and university students are not unusual given that these individuals are at higher risk for psychiatric disorders compared to their non-student counterparts (Blanco et al., 2008; Verger et al., 2010; Eisenberg et al., 2011). Moreover, anxiety disorders are also the most commonly reported psychiatric conditions in undergraduates, with a prevalence of approximately 11.9% (Blanco et al., 2008).

**Collinearity Diagnostics.** Tests were performed to determine if the data on anxiety met the assumption of collinearity (Craney and Surles, 2002; Midi et al., 2010; Fox and Weisberg, 2011). **Table S1** presents the results of these tests, detailing the pre- and post-instruction coefficients and collinearity diagnostics of each dependent variable. For the entire sample, at both time points, for each factor, the variance inflation factor (VIF; $1/(1-R^2)$) did not exceed 2.5, the eigenvalues were greater than zero, and the condition indices ranged from 1 to 2.458. Overall, these results indicate that multicollinearity was not a concern.

### I. Pre-Instruction Collinearity Diagnostics

**Coefficients for Clinical Anxiety as Dependent Variable**

| Model | | Unstandardized Coefficients | | Standardized Coefficients | t | *P*-value | Correlations |
|---|---|---|---|---|---|---|---|
| | | B | Std. Error | Beta | | | Zero-order |
| 1 | (Constant) | 0.002 | 0.091 | | 0.026 | 0.979 | |
| | Spatial Anxiety | -0.042 | 0.100 | -0.042 | -0.423 | 0.673 | 0.136 |
| | Science Anxiety | 0.261 | 0.116 | 0.245 | 2.247 | 0.027 | 0.372 |
| | Math Anxiety | 0.279 | 0.108 | 0.278 | 2.576 | 0.012 | 0.390 |

**Collinearity Diagnostics for Clinical Anxiety as Dependent Variable**

| Model | Dimension | Eigenvalue | Condition Index | Variance Proportions | | | |
|---|---|---|---|---|---|---|---|
| | | | | (Constant) | Spatial | Science | Math |
| 1 | 1 | 1.802 | 1.000 | 0.00 | 0.12 | 0.14 | 0.14 |
| | 2 | 1.001 | 1.341 | 0.99 | 0.00 | 0.00 | 0.00 |
| | 3 | 0.711 | 1.591 | 0.00 | 0.87 | 0.09 | 0.18 |
| | 4 | 0.486 | 1.926 | 0.01 | 0.01 | 0.76 | 0.68 |

**Coefficients for Math Anxiety as Dependent Variable**

| Model | | Unstandardized Coefficients | | Standardized Coefficients | t | *P*-value | Correlations |
|---|---|---|---|---|---|---|---|
| | | B | Std. Error | Beta | | | Zero-order |
| 1 | (Constant) | 0.024 | 0.083 | | 0.287 | 0.775 | |
| | Spatial Anxiety | 0.165 | 0.089 | 0.165 | 1.857 | 0.066 | 0.327 |
| | Science Anxiety | 0.390 | 0.100 | 0.367 | 3.877 | 0.000 | 0.512 |
| | Clinical Anxiety | 0.230 | 0.089 | 0.230 | 2.576 | 0.012 | 0.390 |

**Collinearity Diagnostics for Math Anxiety as Dependent Variable**

| Model | Dimension | Eigenvalue | Condition Index | Variance Proportions | | | |
|---|---|---|---|---|---|---|---|
| | | | | (Constant) | Spatial | Science | Clinical |
| 1 | 1 | 1.588 | 1.000 | 0.00 | 0.15 | 0.21 | 0.16 |
| | 2 | 0.999 | 1.261 | 0.99 | 0.00 | 0.00 | 0.00 |
| | 3 | 0.864 | 1.356 | 0.00 | 0.54 | 0.00 | 0.47 |
| | 4 | 0.548 | 1.703 | 0.00 | 0.31 | 0.79 | 0.37 |

**Coefficients for Spatial Anxiety as Dependent Variable**

| Model | | Unstandardized Coefficients | | Standardized Coefficients | t | *P*-value | Correlations |
|---|---|---|---|---|---|---|---|
| | | B | Std. Error | Beta | | | Zero-order |
| 1 | (Constant) | -0.003 | 0.093 | | -0.033 | 0.974 | |
| | Math Anxiety | 0.208 | 0.112 | 0.208 | 1.857 | 0.066 | 0.327 |
| | Science Anxiety | 0.280 | 0.118 | 0.265 | 2.379 | 0.019 | 0.355 |
| | Clinical Anxiety | -0.044 | 0.103 | -0.044 | -0.423 | 0.673 | 0.136 |

**Collinearity Diagnostics for Spatial Anxiety as Dependent Variable**

| Model | Dimension | Eigenvalue | Condition Index | Variance Proportions | | | |
|---|---|---|---|---|---|---|---|
| | | | | (Constant) | Math | Science | Clinical |
| 1 | 1 | 1.852 | 1.000 | 0.00 | 0.13 | 0.13 | 0.12 |
| | 2 | 1.002 | 1.360 | 0.99 | 0.00 | 0.00 | 0.00 |
| | 3 | 0.660 | 1.676 | 0.00 | 0.12 | 0.18 | 0.87 |
| | 4 | 0.486 | 1.952 | 0.01 | 0.74 | 0.68 | 0.00 |

**Coefficients for Science Anxiety as Dependent Variable**

| Model | | Unstandardized Coefficients | | Standardized Coefficients | t | *P*-value | Correlations |
|---|---|---|---|---|---|---|---|
| | | B | Std. Error | Beta | | | Zero-order |
| 1 | (Constant) | -0.034 | 0.078 | | -0.433 | 0.666 | |
| | Math Anxiety | 0.344 | 0.089 | 0.365 | 3.877 | 0.000 | 0.512 |
| | Spatial Anxiety | 0.197 | 0.083 | 0.208 | 2.379 | 0.019 | 0.355 |
| | Clinical Anxiety | 0.190 | 0.085 | 0.202 | 2.247 | 0.027 | 0.372 |

**Collinearity Diagnostics for Science Anxiety as Dependent Variable**

| Model | Dimension | Eigenvalue | Condition Index | Variance Proportions | | | |
|---|---|---|---|---|---|---|---|
| | | | | (Constant) | Math | Spatial | Clinical |
| 1 | 1 | 1.580 | 1.000 | 0.00 | 0.21 | 0.14 | 0.17 |
| | 2 | 1.001 | 1.257 | 1.00 | 0.00 | 0.00 | 0.00 |
| | 3 | 0.866 | 1.351 | 0.00 | 0.00 | 0.62 | 0.39 |
| | 4 | 0.553 | 1.690 | 0.00 | 0.78 | 0.24 | 0.44 |

### II. Post-Instruction Collinearity Diagnostics

**Coefficients for Clinical Anxiety as Dependent Variable**

| Model | | Unstandardized Coefficients | | Standardized Coefficients | t | *P*-value | Correlations |
|---|---|---|---|---|---|---|---|
| | | B | Std. Error | Beta | | | Zero-order |
| 1 | (Constant) | 0.000 | 0.094 | | -0.001 | 0.999 | |
| | Spatial Anxiety | -0.084 | 0.099 | -0.084 | -0.853 | 0.396 | 0.024 |
| | Science Anxiety | 0.268 | 0.140 | 0.253 | 1.912 | 0.059 | 0.341 |
| | Math Anxiety | 0.169 | 0.131 | 0.166 | 1.287 | 0.201 | 0.321 |

**Collinearity Diagnostics for Clinical Anxiety as Dependent Variable**

| Model | Dimension | Eigenvalue | Condition Index | Variance Proportions | | | |
|---|---|---|---|---|---|---|---|
| | | | | (Constant) | Spatial | Science | Math |
| 1 | 1 | 1.834 | 1.000 | 0.00 | 0.08 | 0.12 | 0.12 |
| | 2 | 0.998 | 1.356 | 1.00 | 0.00 | 0.00 | 0.00 |
| | 3 | 0.857 | 1.463 | 0.00 | 0.89 | 0.02 | 0.08 |
| | 4 | 0.311 | 2.428 | 0.00 | 0.04 | 0.86 | 0.80 |

**Coefficients for Math Anxiety as Dependent Variable**

| Model | | Unstandardized Coefficients | | Standardized Coefficients | t | *P*-value | Correlations |
|---|---|---|---|---|---|---|---|
| | | B | Std. Error | Beta | | | Zero-order |
| 1 | (Constant) | 0.003 | 0.072 | | 0.042 | 0.966 | |
| | Spatial Anxiety | 0.009 | 0.076 | 0.009 | 0.116 | 0.908 | 0.202 |
| | Science Anxiety | 0.672 | 0.086 | 0.644 | 7.823 | 0.000 | 0.681 |
| | Clinical Anxiety | 0.100 | 0.077 | 0.101 | 1.287 | 0.201 | 0.321 |

**Collinearity Diagnostics for Math Anxiety as Dependent Variable**

| Model | Dimension | Eigenvalue | Condition Index | Variance Proportions | | | |
|---|---|---|---|---|---|---|---|
| | | | | (Constant) | Spatial | Science | Clinical |
| 1 | 1 | 1.467 | 1.000 | 0.00 | 0.14 | 0.26 | 0.17 |
| | 2 | 0.997 | 1.213 | 0.98 | 0.00 | 0.00 | 0.02 |
| | 3 | 0.976 | 1.226 | 0.01 | 0.53 | 0.00 | 0.37 |
| | 4 | 0.560 | 1.619 | 0.00 | 0.33 | 0.73 | 0.44 |

**Coefficients for Spatial Anxiety as Dependent Variable**

| Model | | Unstandardized Coefficients | | Standardized Coefficients | t | *P*-value | Correlations |
|---|---|---|---|---|---|---|---|
| | | B | Std. Error | Beta | | | Zero-order |
| 1 | (Constant) | -0.003 | 0.096 | | -0.033 | 0.974 | |
| | Math Anxiety | 0.016 | 0.135 | 0.015 | 0.116 | 0.908 | 0.202 |
| | Science Anxiety | 0.334 | 0.142 | 0.316 | 2.354 | 0.021 | 0.296 |
| | Clinical Anxiety | -0.088 | 0.103 | -0.088 | -0.853 | 0.396 | 0.024 |

**Collinearity Diagnostics for Spatial Anxiety as Dependent Variable**

| Model | Dimension | Eigenvalue | Condition Index | Variance Proportions | | | |
|---|---|---|---|---|---|---|---|
| | | | | (Constant) | Math | Science | Clinical |
| 1 | 1 | 1.922 | 1.000 | 0.00 | 0.11 | 0.11 | 0.09 |
| | 2 | 0.998 | 1.387 | 1.00 | 0.00 | 0.00 | 0.00 |
| | 3 | 0.762 | 1.588 | 0.00 | 0.08 | 0.06 | 0.90 |
| | 4 | 0.318 | 2.458 | 0.00 | 0.81 | 0.83 | 0.00 |

**Coefficients for Science Anxiety as Dependent Variable**

| Model | | Unstandardized Coefficients | | Standardized Coefficients | t | *P*-value | Correlations |
|---|---|---|---|---|---|---|---|
| | | B | Std. Error | Beta | | | Zero-order |
| 1 | (Constant) | -0.019 | 0.067 | | -0.278 | 0.782 | |
| | Math Anxiety | 0.576 | 0.074 | 0.601 | 7.823 | 0.000 | 0.681 |
| | Spatial Anxiety | 0.162 | 0.069 | 0.171 | 2.354 | 0.021 | 0.296 |
| | Clinical Anxiety | 0.136 | 0.071 | 0.144 | 1.912 | 0.059 | 0.341 |

**Collinearity Diagnostics for Science Anxiety as Dependent Variable**

| Model | Dimension | Eigenvalue | Condition Index | Variance Proportions | | | |
|---|---|---|---|---|---|---|---|
| | | | | (Constant) | Math | Spatial | Clinical |
| 1 | 1 | 1.393 | 1.000 | 0.00 | 0.30 | 0.11 | 0.23 |
| | 2 | 0.999 | 1.181 | 0.94 | 0.00 | 0.03 | 0.03 |
| | 3 | 0.977 | 1.194 | 0.05 | 0.00 | 0.68 | 0.23 |
| | 4 | 0.631 | 1.486 | 0.00 | 0.70 | 0.19 | 0.51 |

**Table S1.** Collinearity Diagnostics. *Pre- and post-instruction coefficients and collinearity diagnostics for clinical, math, spatial, and science anxiety. The* Dimension *column denotes the linear combination of the* *four dependent variables in order of largest to smallest possible variance combination. The* Eigenvalue *column denotes the variance of each linear combination. Eigenvalues close to 0 indicate that the factors are highly intercorrelated. The* Condition Index *column denotes the calculation of the square roots of the ratios of the largest eigenvalue to each successive eigenvalue, as computed by SPSS. The* Variance Proportions *column is utilized to assess how each dimensions' variance proportions are related to one another. For example, in the table, Collinearity Diagnostics for Science Anxiety as Dependent Variable, 68% of the variance for Spatial Anxiety is associated with Dimension 3 (Clinical Anxiety), which has an eigenvalue of 0.977 and a condition index of 1.194. Collinearity is problematic if two or more variables have variance proportions of 0.50 or more that correspond to condition indices of 30 or larger (Osborne, 2000; Pedhazur, 1997).*

**Sex Differences in STEM Anxiety.** We performed mixed model ANOVA analyses for each anxiety measure (**Table 1**). These analyses demonstrated significant main effects of sex on all measures of anxiety, including science, spatial, math, and clinical anxiety. To assess the robustness of these results to potential violations of the assumptions of normality and equal variances, we replicated the analyses using robust ANOVA methods recommended by Wilcox (2017). The robust ANOVAs returned the same pattern of results as the classical ANOVAs, strengthening our confidence in these findings. Additionally, we note that the same general pattern of results held when running the analyses using linear mixed model (multilevel) regressions, both with and without controlling for clinical anxiety when analyzing the remaining anxiety measures (**Table S2**).

| | Not Controlling for Clinical Anxiety | | | | Controlling for Clinical Anxiety | | |
|---|---|---|---|---|---|---|---|
| Coefficient | Spatial Anxiety | Science Anxiety | Math Anxiety | Clinical Anxiety | Spatial Anxiety | Science Anxiety | Math Anxiety |
| Intercept | 6.59*** | 3.15** | 17.67*** | 6.02*** | 6.50*** | 1.76 | 15.42*** |
| | (0.57) | (1.12) | (2.37) | (1.26) | (0.60) | (1.13) | (2.37) |
| Time | -0.48 | 8.13*** | 1.60 | 1.40 | -0.50 | 7.81*** | 1.08 |
| | (0.56) | (1.22) | (2.03) | (1.48) | (0.56) | (1.22) | (2.06) |
| Gender | 1.93* | 3.25$^{+}$ | 11.84** | 4.70* | 1.86* | 2.17 | 10.08** |
| | (0.85) | (1.66) | (3.51) | (1.86) | (0.86) | (1.61) | (3.40) |
| Time*Gender | 0.72 | 1.89 | -1.34 | -2.38 | 0.75 | 2.44 | -0.45 |
| | (0.82) | (1.80) | (3.01) | (2.20) | (0.83) | (1.81) | (3.05) |
| Clinical Anxiety | -- | -- | -- | -- | 0.02 | 0.23*** | 0.37** |
| | | | | | (0.03) | (0.06) | (0.11) |

**Table S2.** Parameter Estimates (Standard Errors) from Mixed Model Analyses of Anxiety Measures. *Time is coded as 0 = pre-instruction, 1 = post-instruction. Sex is coded as 0 = male, 1 = female.* $^{+}$ *indicates p = 0.05, * indicates p < 0.05, ** indicates p < 0.05, *** indicates p < 0.001.*

**Correlations between Changes in Anxiety and Changes in Connectivity.** **Fig. 3** displays the observed correlations between anxiety (e.g., science, spatial, math, and clinical) and inter-network functional connectivity (e.g., CEN-DMN, DMN-SN, and SN-CEN) at pre-instruction, post-instruction, and the change across time. **Fig. S2** provides additional scatterplots to aid in the interpretation of correlations between the change in anxiety scores and the change in connectivity from pre- to post-instruction (i.e., bottom of **Fig. 3**).

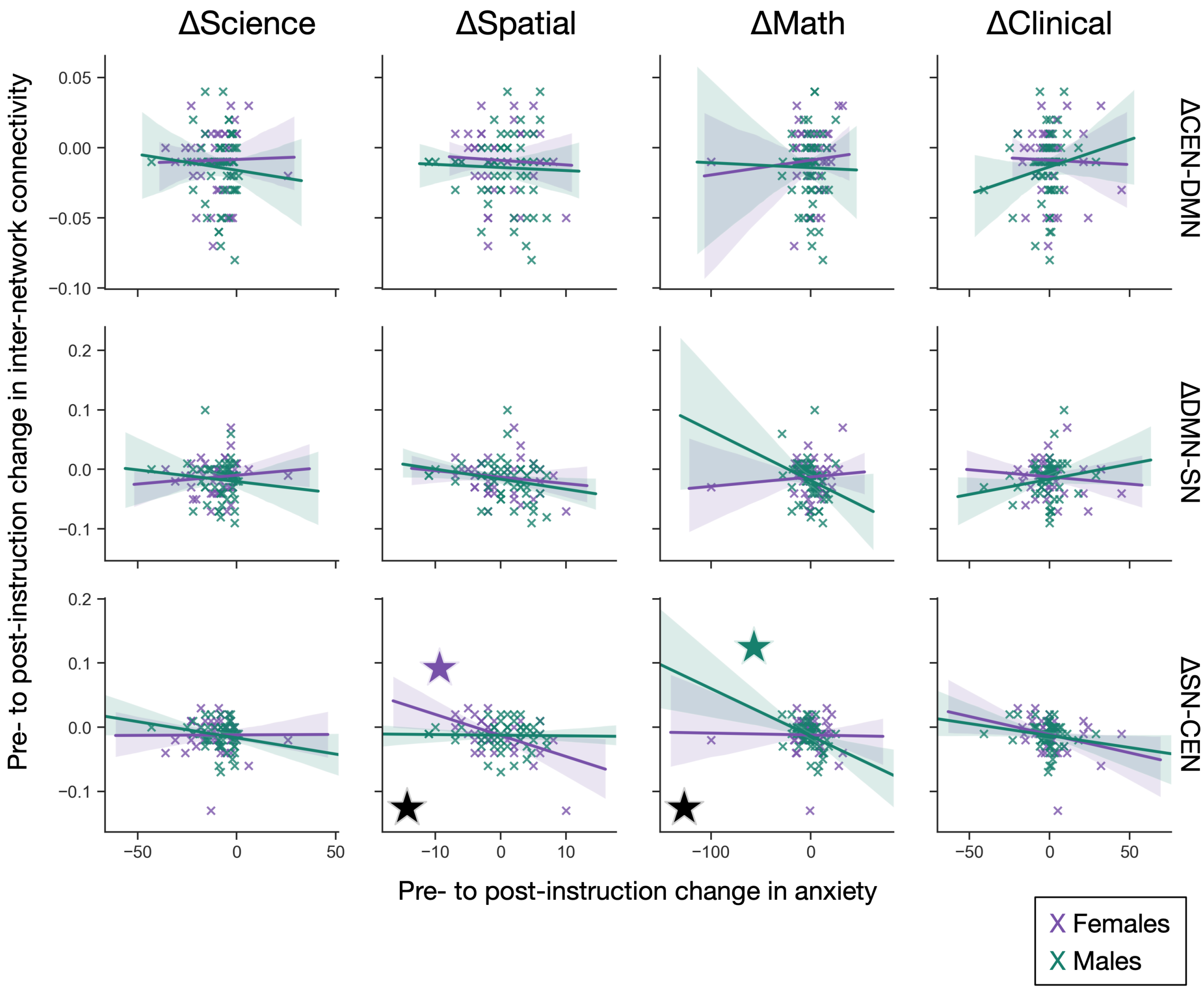


**Fig. S2.** Changes Across Time in Anxiety and Connectivity. *Reflecting the "POST > PRE" panel of Fig. 3, scatterplots are provided to detail each participant's pre- to post-instruction change in functional connectivity, per network pair, against their change in anxiety score. Significant within-sex correlations between these changes are indicated by either a purple or green star, depending on whether the correlation was significant for female or male students, respectively. Significant between-sex differences in these correlations are indicated by a black star.*

**Additional Analyses: Controlling for Clinical Anxiety.** Although our primary analyses treated STEM (science, spatial, and math) and clinical anxiety as independent constructs, we acknowledge that this may not be the case for some students. To assess the relationships between STEM anxiety and functional connectivity, while controlling for the effect of clinical anxiety, we conducted additional analyses of the data by computing partial Pearson correlations. Among female students at pre-instruction, when controlling for clinical anxiety, no significant partial correlations were observed between anxiety scores and inter-network connectivity. In contrast, when controlling for clinical anxiety, male students at pre-instruction exhibited significant partial correlations between science anxiety and the DMN-SN inter-network connectivity ($r(55) = 0.317$, $P = 0.020$), spatial anxiety and CEN-DMN ($r = 0.352$, $P = 0.009$), math anxiety and CEN-DMN ($r = 0.284$, $P = 0.038$), and math anxiety and DMN-SN ($r = 0.374$, $P = 0.005$). Among female students at post-instruction, when controlling for clinical anxiety, no significant partial correlations were observed between anxiety scores and inter-network connectivity. In contrast, when controlling for clinical anxiety, male students at post-instruction exhibited significant partial correlations between spatial anxiety and CEN-DMN connectivity ($r(55) = 0.383$, $P = 0.004$), spatial anxiety and DMN-SN ($r = 0.437$, $P = 0.001$), and math anxiety and DMN-SN ($r = 0.340$, $P = 0.012$). Importantly, these supplemental results are consistent with our primary results shown in **Fig. 3**, producing approximately equal, and even in some instances stronger, associations between STEM anxiety and functional connectivity in male students when controlling for clinical anxiety.

Next, we examined the correlations between anxiety scores and academic performance, controlling for clinical anxiety. Among female students at pre-instruction, when controlling for clinical anxiety, GPA was positively partially correlated with spatial anxiety ($r(44) = 0.413$, $P = 0.006$). Among male students at pre-instruction, when controlling for clinical anxiety, GPA was negatively partially correlated with math anxiety ($r(49) = -0.330$, $P = 0.022$). Among female students at post-instruction, when controlling for clinical anxiety, no significant partial correlations were observed between anxiety and academic performance. Among male students at post-instruction, when controlling for clinical anxiety, GPA was negatively partially correlated with science anxiety ($r(49) = -0.360$, $P = 0.012$) and math anxiety ($r(49) = -0.449$, $P = 0.001$), grade and science anxiety ($r(55) = -0.341$, $P = 0.012$), and grade was negatively partially correlated with math anxiety ($r(55) = -0.415$, $P = 0.002$). Again, these supplemental results are generally consistent with our primary results shown in **Fig. 4**, although some significant correlations in female students were no longer observed to be significant after controlling for clinical anxiety.

Lastly, we investigated if rs-fMRI connectivity was correlated with academic performance at pre- or post-instruction, controlling for clinical anxiety. Among female students, when controlling for clinical anxiety, no significant partial correlations were observed between inter-network brain connectivity and GPA or course grade at either time point. Among male students at post-instruction, when controlling for clinical anxiety, DMN-SN connectivity was negatively partially correlated with course grade ($r(55) = -0.296$, $P = 0.030$) and SN-CEN connectivity was negatively partially correlated with course grade ($r(55) = -0.271$, $P = 0.047$). These supplemental results are consistent with our primary results shown in **Fig. 5a** and provide added insight to the relationship between SN-CEN and academic performance. Overall, our results when controlling for clinical anxiety provide strong support for the primary results presented in this study.

### SUPPLEMENTAL INFORMATION REFERENCES


Beck, A. T., Epstein, N., Brown, G. & Steer, R. A. An inventory for measuring clinical anxiety: Psychometric properties. Journal of Consulting and Clinical Psychology 56(6), 893–897 (1988). doi:10.1037/0022-006X.56.6.893.

Betancourt, H. & López, S. R. The study of culture, ethnicity, and race in american psychology. *American Psychologist* **48,** 629–637 (1993).

Bilogur, A. Missingno: A missing data visualization suite. Journal of Open Source Software, 3(22), 547 (2018). https://doi.org/10.21105/joss.00547

Blanco, C. *et al.* Mental health of college students and their non-college-attending peers: Results from the national epidemiologic study on alcohol and related conditions. *Archives of General Psychiatry* **65,** 1429–1437 (2008).

Craney, T. A. & Surles, J. G. Model-dependent variance inflation factor cutoff values. *Quality Engineering* **14,** 391–403 (2002).

Eisenberg, D., Hunt, J., Speer, N. & Zivin, K. Mental health service utilization among college students in the United States. *Journal of Nervous and Mental Disease* **199,** 301–308 (2011).

Fox, J. & Weisberg, S. in *An R companion to applied regression* 285–328 (Sage, 2011).

Midi, H., Sarkar, S. K. & Rana, S. Collinearity diagnostics of binary logistic regression model. *Journal of Interdisciplinary Mathematics* **13,** 253–267 (2010).

Oldfield, R. C. The assessment and analysis of handedness: The Edinburgh inventory *Neuropsychologia* 9(1), 97–113 (1971). doi:10.1016/0028-3932(71)90067-4.

Osborne, J. W. Prediction in Multiple Regression. Practical Assessment, Research & Evaluation **7,** 1–6 (2000).

Pedhazur, E. J. (1997). Multiple regression in behavioral research (3rd ed.). Orlando, FL: Harcourt Brace.

Phinney, J. S. When We Talk about American Ethnic Groups, What Do We Mean? *American Psychologist* **51,** 918–927 (1996).

U.S. Census Bureau (2017a). *Research to Improve Data on Race and Ethnicity.* Retrieved from https://www.census.gov/about/our-research/race-ethnicity.html

U.S. Census Bureau (2017b). *Race & Ethnicity.* Retrieved from https://www.census.gov/mso/www/training/pdf/race-ethnicity-onepager.pdf

Verger, P., Guagliardo, V., Gilbert, F., Rouillon, F. & Kovess-Masfety, V. Psychiatric disorders in students in six French universities: 12-month prevalence, comorbidity, impairment and help-seeking. *Social Psychiatry and Psychiatric Epidemiology* **45,** 189–199 (2010).

Wilcox, R. (2017). Modern statistics for the social and behavioral sciences: A practical introduction. (2nd ed.). Boca Raton, FL: CRC Press.
